## Supplemental Files for "*arfA* antisense RNA regulates MscL excretory activity"

Rosa Morra *et al.*

**This file includes:**

**Figure S1. Taxonomic analysis of bacteria in which only *arfA*, *mscL* or neither of them are present.**

**Figure S2. Design of *arfA* and *mscL* deleted (Δ*arfA*) strains**

**Figure S3. Growth induced hypoosmotic stress**

**Figure S4. Analysis of *mscL* RNA and recombinant protein localisation in *E. coli* BL21(DE3)::*mscLHis***

**Figure S5. Analysis of *arfA* RNA**

**Figure S6. The predicted secondary strucure of *mscL* 3' UTR and *arfA* 3' CDS**

**Figure S7. RNase III in vitro assay**

**Table S1 Enrichment analysis of genomic cluster across taxa groups**

**Table S2. Syntenic analysis of *arfA* proximal and overlapping genes**

**Table S3. *E. coli* strains and plasmids used in this study**

**Table S4. Primers used for cloning and generation of targeted gene deletion strains (source IDT)**

**Table S5. Primers used for qRT-PCR (source IDT)**

**Data S1. (separate file)**

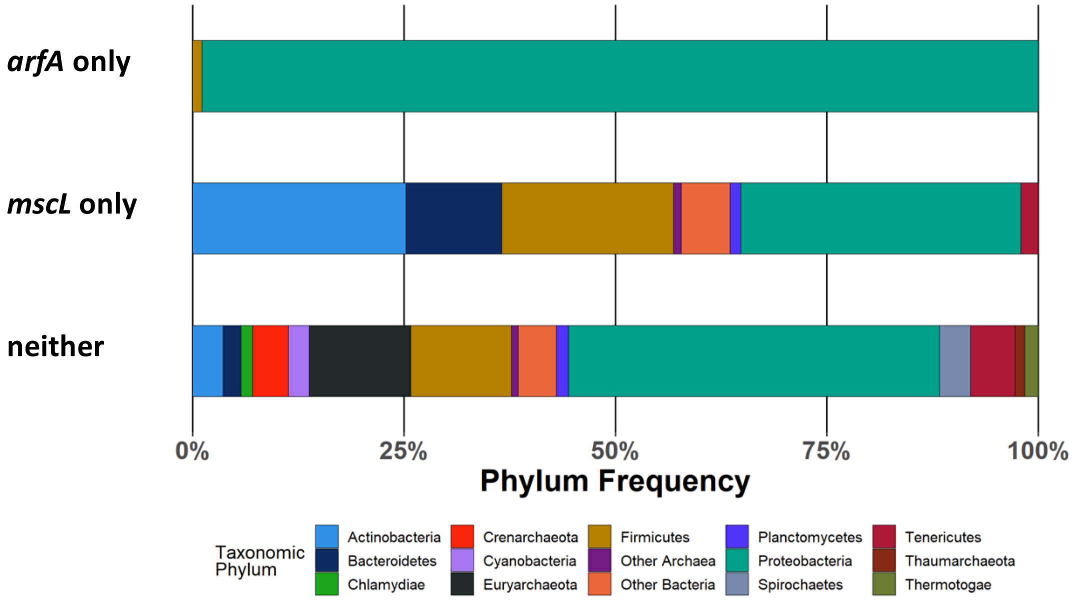


**Figure S1. Taxonomic analysis of bacteria in which only *arfA*, *mscL* or neither of them are present.** See Data S1 for genomic and taxonomic data.

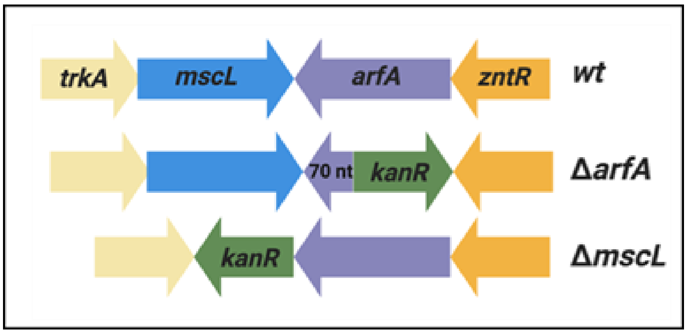


**Figure S2. Design of *arfA* and *mscL* deleted (Δ*arfA*) strains.** The 3′ regon of *arfA* CDS (70 nucleotides) was maintained in Δ*arfA* strains, as the genomic region is a part of *mscL* 3′ UTR. The kanamycin selection cassette (*kan*R) was inserted in the opposite orientation to the target gene to avoid read-through transcription. The direction of transcription of genes is indicated by the arrows.

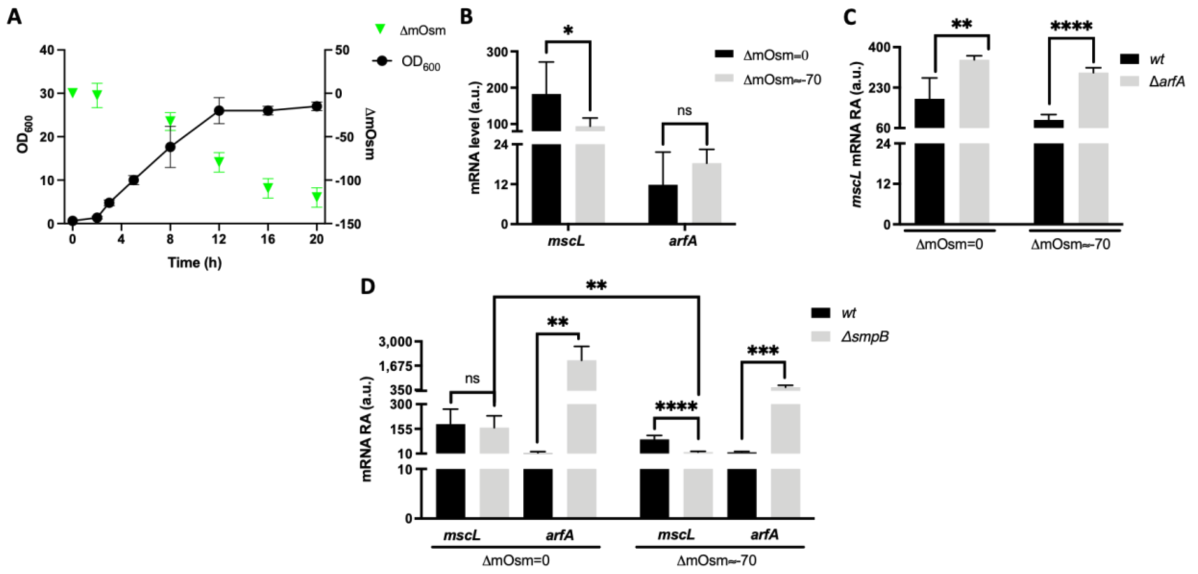


**Figure S3**. **Effect of media osmolality and trans-translation upon *mscL* and *arfA* expression (A)** Representative *E. coli* growth curve at 30°C in rich media (TB) and the decline in media osmolality (ΔmOsm) caused by the cell growth ($OD_{600}$). Relative *mscL* and *arfA* RNA abundance (RA) measured in strains grown in TB media pre (ΔmOsm=0, OD600 ≈ 4) and post (ΔmOsm~70, OD600 ≈ 12) drop in osmolality for **(B)** *wt,* **(C)** *wt* and *arfA* deleted (Δ*arfA*), and **(D)** *wt* and *smpB* deleted (Δ*smpB*) *E. coli* K12 strains. Ct values were normalised to the RNA abundance of the most stable housekeeping genes of *E. coli*. The error bars represent the standard deviation of at least three biological replicates; unpaired t-test analysis was performed between pre vs. post osmolality drop **(B)** and *wt* vs gene-specific deleted strain **(C, D)** (P < 0.05*, < 0.01**, < 0.001***, < 0.0001****). ns = no significance. a.u. = arbitrary units.

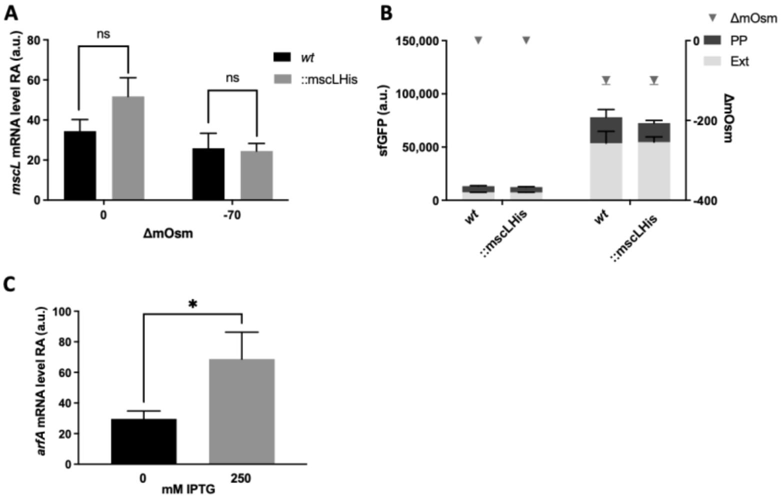


**Figure S4. Analysis of *mscL* RNA and recombinant protein localisation in *E. coli* BL21(DE3)::*mscLHis*. (A)** Relative *mscL* abundance (RA) and localization of recombinant sfGFP **(B)** detected in *E. coli* BL21(DE3) *wt* and ::*mscLHis* cells grown in TB media pre (ΔmOsm=0, $OD_{600} \approx 4$) and post (ΔmOsm~70, OD600 ≈ 12) drop in media osmolality. PP = periplasmic fraction; Ext = extracellular fraction. **(C)** Relative *arfA* abundance (RA) measured in *E. coli* BL21(DE3)::*mscLHis* cells grown in TB media post drop in osmolality (ΔmOsm~70, $OD_{600} \approx 12$) in absence or presence of IPTG. Unpaired t-test analysis was performed in **(A)** and **(C).** ($P < 0.05$*). a.u = arbitrary units.

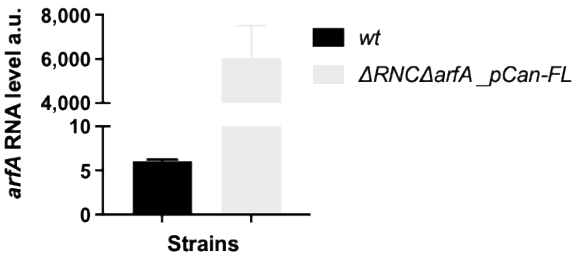


**Figure S5. Analysis of *arfA* RNA.** Native and episomal *arfA* relative abundance (RA) measured in *wt* and Δ*RNC*Δ*arfA E. coli* cells, *,* bearing an empty and pCan-FL palsmid respectively*,* grown in LB media.

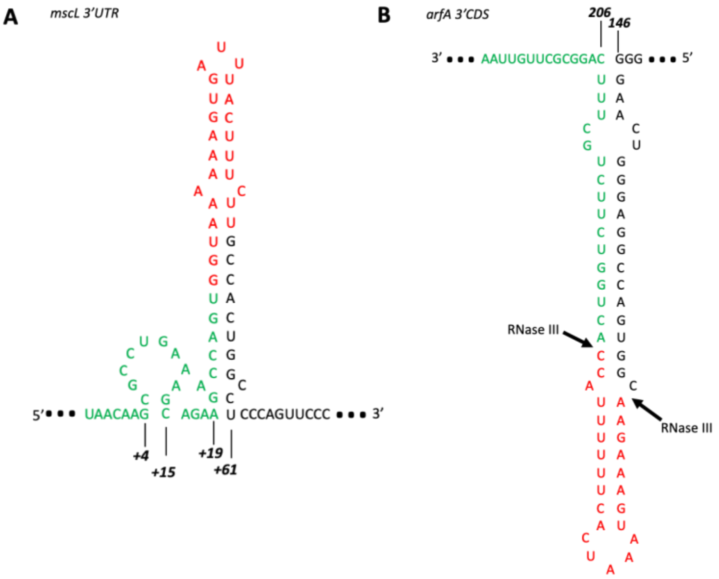


**Figure S6. The predicted secondary strucure of *mscL* 3' UTR and *arfA* 3' CDS. (A)** The *mscL* 3' UTR stem loop seconday structure was predicted using the RNA Vienna RNAfold web server submitting the *mscL* 3' UTR sequence between the stop codon (UAA) and +70 nucleotide. **(B)** The *arfA* 3' CDS stam loop seconday structure was predicted using the RNA Vienna RNAfold web server submitting the *arfA* CDS sequence between nucleodite 144 ad stop codon (UAA). The complemetary region between *arfA* sRNA relased by the cleavge of RNAseIII and *mscL* 3' UTR is shown in red and green.

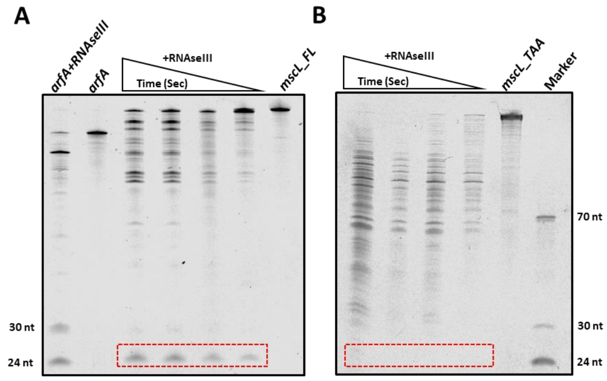


**Figure S7. RNase III in vitro assay.** RNaseIII cleavage assay with the *in vitro* synthesized transcripts representing the **(A)** full length transcript (*mscL_FL*) including the 3' UTR, and **B)** a truncated CDS only transcript (*mscL_TAA*). Cleavage reactions were incubated for 7.5, 15, 30 and 60 sec at 37°C. The red dashed box highlights the ~25nt sRNA released from *mscL_FL,* but not from *mscL_TAA*. *arfA* transcript (*arfA*) which releases 24 and 30 nt sRNAs after cleavage by RNAseIII, was used as positive control.

**Table S1. Enrichment analysis of genomic cluster across taxa groups**

| Phyla/Class | Cluster | Cluster Count | Total Count | Percentage |
|---|---|---|---|---|
| *Actinobacteria* | mscl_only | 584 | 625 | 93.4 |
| *Firmicutes* | mscl_only | 470 | 608 | 77.3 |
| *Alphaproteobacteria* | mscl_only | 321 | 422 | 76.1 |
| *Bacteroidetes* | mscl_only | 262 | 286 | 91.6 |
| *Gammaproteobacteria* | no_proteins | 237 | 763 | 31.1 |
| *Betaproteobacteria* | mscl_only | 213 | 271 | 78.6 |
| *Gammaproteobacteria* | mscl_only | 173 | 763 | 22.7 |
| *Gammaproteobacteria* | distal | 137 | 763 | 18.0 |
| *Euryarchaeota* | no_proteins | 136 | 153 | 88.9 |
| *Firmicutes* | no_proteins | 135 | 608 | 22.2 |
| *Gammaproteobacteria* | proximal | 103 | 763 | 13.5 |
| *Alphaproteobacteria* | no_proteins | 100 | 422 | 23.7 |
| *Gammaproteobacteria* | arfa_only | 88 | 763 | 11.5 |
| *Epsilonproteobacteria* | no_proteins | 66 | 94 | 70.2 |
| *Tenericutes* | no_proteins | 60 | 108 | 55.6 |
| *Deltaproteobacteria* | no_proteins | 50 | 75 | 66.7 |
| *Tenericutes* | mscl_only | 48 | 108 | 44.4 |
| *Crenarchaeota* | no_proteins | 48 | 48 | 100.0 |
| *Actinobacteria* | no_proteins | 41 | 625 | 6.6 |
| *Spirochaetes* | no_proteins | 41 | 48 | 85.4 |

Note: Subset of taxonomic groups shown, only those containing >1% of total number of genomes present. See Data S1 for genomic and taxonomic data.

### Table S2 Syntenic analysis of *arfA* proximal and overlapping genes

| NCBI protein annotation | Total count | Average ArfA Bit score ± SD | Genera | ArfA-only count | Distal count |
|---|---|---|---|---|---|
| iron-sulfur cluster assembly protein IscA | 10 | 75.7 ± 2.7 | *Neisseria* | 4 | 6 |
| threonine/serine exporter family protein | 9 | 84.1± 2.5 | *Vibrio* | 9 | 0 |
| 30S ribosomal protein S12 methylthiotransferase RimO | 9 | 83.0 ± 4.2 | *Shewanella* | 9 | 0 |
| hypothetical protein | 6 | 70.8 ± 22.5 | *multiple* | 3 | 3 |
| Trk system potassium transporter TrkA | 4 | 89.5 ± 2.2 | *Xenorhabdus* | 4 | 0 |
| dethiobiotin synthase | 4 | 77.8 ± 3.8 | *Mannheimia* | 2 | 2 |
| cytochrome b | 3 | 73.3 ± 0.3 | *Pseudoalteromonas* | 3 | 0 |
| radical SAM family heme chaperone HemW | 3 | 83.2 ± 4.1 | *Aeromonas* | 0 | 3 |
| diguanylate cyclase | 2 | 84.9 ± 2.1 | *Shewanella*, *Pseudoalteromonas* | 2 | 0 |
| pyrimidine 5'-nucleotidase | 2 | 82.5 ± 0 | *Agarivorans* | 2 | 0 |
| DUF3465 domain-containing protein | 2 | 82.1 ± 1.6 | *Shewanella* | 2 | 0 |
| helix-turn-helix domain-containing protein | 2 | 88.7 ± 6.7 | *Leminorella*, *Pragia* | 0 | 2 |
| oxygen-dependent coproporphyrinogen oxidase | 1 | 73 | *Colwellia* | 1 | 0 |
| FNR family transcription factor | 1 | 82.2 | *Frederiksenia* | 0 | 1 |
| amino acid ABC transporter substrate-binding protein | 1 | 85.8 | *Shewanella* | 1 | 0 |
| YqaE/Pmp3 family membrane protein | 1 | 78.7 | *Dongshaea* | 1 | 0 |
| response regulator transcription factor | 1 | 53 | *Halomonas* | 0 | 1 |
| PAS domain-containing methyl-accepting chemotaxis protein | 1 | 83.8 | *Vibrio* | 1 | 0 |
| diacylglycerol kinase | 1 | 80.8 | *Vibrio* | 1 | 0 |
| recombinase RecA | 1 | 78.9 | *Paralysiella* | 0 | 1 |
| sulfate transporter CysZ | 1 | 87.6 | *Haemophilus* | 0 | 1 |
| NCS2 family permease | 1 | 35.1 | *Cardiobacterium* | 0 | 1 |
| methyltransferase domain-containing protein | 1 | 78.2 | *Pasteurella* | 0 | 1 |
| methylenetetrahydrofolate reductase | 1 | 75.1 | *Bibersteinia* | 0 | 1 |
| N-acetyltransferase | 1 | 85 | *Plesiomonas* | 0 | 1 |
| IS982 family transposase | 1 | 88.7 | *Xenorhabdus* | 1 | 0 |
| LacI family DNA-binding transcriptional regulator | 1 | 73 | *Pseudoalteromonas* | 1 | 0 |

Genes located within 110 nucleotides from *arfA* were included. See Data S1 for genomic and taxonomic data.

**Table S3.** E. coli strains and plasmids used in this study.

| Strain or Plasmid | Description | Reference or source |
|---|---|---|
| **Strain** | | |
| BL21 (DE3) | | |
| • Wild-type | | • Novagen |
| • Wild-type *mscLHis* | • Replacement of wild-type *mscL* with *mscLHis* gene tagged with hexahistidine at C-terminus. Containing kanamycin resistance cassette | • This study |
| • Δ*arfA mscLHis* | • A knockout mutant of *arfA*. Deletion of transcription starting site (TSS) and first 43 amino acids of ArfA (1-43aa). Replacement of wild-type *mscL* with *mscLHis* gene tagged with hexahistidine at C-terminus. Containing kanamycin resistance cassette | • This study |
| K-12 MG1655 | | |
| • Wild-type | | • Lab stock |
| • Δ*arfA* | • A knockout mutant of *arfA*. Deletion of transcription starting site (TSS) and first 43 amino acids ArfA (1-43aa). Containing kanamycin resistance cassette. | • This study |
| • Δ*mscL* | • A knockout mutant of *mscL*. Deletion of the promoter, 5'UTR and CDS of *mscL*. Containing kanamycin resistance cassette. | • This study |
| K-12 BW25113 | | |
| • Δ*rpoS* | • A knockout mutant of *rpoS*. Containing kanamycin resistance cassette | • Keio collection (1) |
| • Δ*smpB* | • A knockout mutant of *smpB*. Containing kanamycin resistance cassette | |
| MC1061 | | |
| • Δ*rnc* | • A non-functional mutant of RNAseIII (Δ*rnc*-38) Containing kanamycin resistance cassette | • (2) |
| **Plasmid** | | |

| Strain or Plasmid | Description | Reference or source |
|---|---|---|
| p131B | | • Lab stock |
| • p131B_P$_{mscL}$-sfGFP | • p131B plasmid cloned with *mscL* promoter fused with *sfGFP* reporter (P$_{mscL}$-sfGFP) as a construct to study promoter activity. Ampicillin or carbenicillin resistance | • This study |
| • p131B_P$_{arfA}$-sfGFP | • p131B plasmid cloned with *arfA* promoter fused with *sfGFP* reporter (P$_{arfA}$-sfGFP) as a construct to study promoter activity. Ampicillin or carbenicillin resistance | • This study |
| pET44 | | • Novagen |
| • pET44-sfGFP | • pET44 (expression plasmid) cloned with *sfGFP* for recombinant protein overexpression (IPTG inducible) and monitoring recombinant protein excretion during osmotic and translational stress conditions. Ampicillin or carbenicillin resistance | • This study |
| pSIM18 | • Expressing λ-red recombinase for homologous recombination of target DNA into *E. coli* genome. Used to integrate (recombination) Kan resistance cassette for knockout mutation of *arfA* (Δ*arfA*) and *mscL* (Δ*mscL*) in K-12 MG1655, and to integrate *mscLHis*-Kan resistance cassette for *mscL* replacement with *mscLHis* (wild-type) and along with knockout mutation of arfA (Δ*arfA*) in BL21 (DE3). Hygromycin resistance | • A gift from SynBioChem, University of Manchester (3) |
| pRL128 | • Used to generate Kan-resistance cassette for chromosome engineering (knockout and gene replacement) | • Lab stock |

| Strain or Plasmid | Description | Reference or source |
|---|---|---|
| pCA24N | • Expression plasmid. T5-based promoter for recombinant gene expression (IPTG inducible). Gentamycin resistance. Used to generate rescue strain (plasmid expression) for gene expression study | • Lab stock |
| • p*FL* | • CDS of *arfA* gene cloned in pCA24N | • This study |
| • p*Δ(154-216nt)* | • CDS of *arfA* gene missing fragment from nucleotide 154 o 216 cloned in pCA24N | • This study |
| • p*IL* | • CDS of *arfA* gene containing inverted repeats at nucleotide position 147 (G into A), 156 (G into A), 159 (C into A), 162 (T into C) and 171 (A into G) cloned in pCA24N | • This study |
| • p*A18T* | • CDS of *arfA* gene containing the substitution of the amino acid Alanine (AGC) in position 18 with a Threonine (ACC) cloned in pCA24N | • This study |
| • p*sRNA* | • Fragment from nucleotide 160 to 203 from *arfA* CDS gene cloned in pCA24N | • This study |

**Table S4. Primers used for cloning and generation of targeted gene deletion strains (source IDT)**

| Primer set | Sequence | Use |
|---|---|---|
| arfA_Δ*(154-216nt)* F<br>+<br>arfA_Δ (154-216nt) R | TAACTATGCGGCCGCTAAGGG<br><br>P-CCAGTTCCCCCGATTGCC | Used to generate p*Δ(154-216nt)* by PCR mutagenesis of p*FL* |
| H1_Δ(P-44) *mscL*<br>+<br>H2_*mscL* | CTAATGACGCCTTATTATTTCCCTTTGATTATCAAGGATT AATTAAATTC*GAGGTGTAGGCTGGAGCTGC*<br>ACCACTGGTCTTCTGCTTTCAGGCGCTTGTTAAGAGCGG TTATTCTGCT*CTTGGGATCCGTCGACCTGCA* | Used to amplify the FRT-flanked kanamycin selection cassettes by PCR to generate Δ*mscL* strains |
| H2_Δ (TSS)*arfA*<br>+<br>H1_*arfA* | CTTGAACAAGGGGCGAGTGGCGTTAAGAGTGGTTGTTG ATTTTTTGCACT*GAGGTGTAGGCTGGAGCTGC*<br>ACCAGTGGTAAAAAAGTGATTTACTTTCTTGCCACTGGC CTCCCAGTTC*C TTGGGATCCGTCGACCTGCA* | Used to amplify the FRT-flanked kanamycin selection cassettes by PCR to generate Δ*arfA* strains |
| H1_Δ(P-44) *mscL* V2 | CTAATGACGCCTTATTATTTCCCTTTGATTATCAAGGATT AATTAAATTC*TTGGGATCCGTCGACCTGCA* | Used to amplify, with H2_Δ (TSS)*arfA*, FRT-flanked kanamycin selection cassettes by PCR to generate Δ*mscL*Δ*arfA* strains |
| H1_*smpB*<br>+<br>H2_*smpB* | GATATGGGGTGTTTTCGATTTCAGATTACCGATGATTCA CGACGCTTATG*TTGGGATCCGTCGACCTGCA*<br>AGGAACTGGTCAATAATTGGAGTGCAGGTTTAACGGTG GGCGTTTTTCAT*GAGGTGTAGGCTGGAGCTGC* | Used to amplify FRT-flanked kanamycin selection cassettes by PCR to generate Δ*smpB* strain |

**Table S5**. Primers used for qRT-PCR (source IDT)

| Primer name | Sequence | Use<br>amplicon size (nt) | %E (F+R) | E (F+R) | Slope |
|---|---|---|---|---|---|
| *msc*L F | TGCCTCCTCTGGGCTTATTA | *mscL* transcript quantification<br><br>96 | 110 | 2.111 | -3.08 |
| *msc*L R | GCATCACAACAGCAGGGATA | Locus_Tag b3291 | | | |
| *arf*A F | TCAGCATACTAAAGGGCAGATAAA | *arfA* transcript quantification<br>76 | 98 | 1.939 | -3.38 |
| *arf*A R | CTACGCGCTGTCGGAATAA | Locus_Tag b4550 | | | |
| *rec*A F | TACCGGTTCGCTTTCACTG | *recA* transcripts quantification<br>76 | 99 | 2.038 | -3.23 |
| *rec*A R | TCCGTAGATTTCGACGATACG | Locus_Tag b2699 | | | |
| *cys*G F | CACTGCGTACCCCAGGAA | *cysG* transcripts quantification (4)<br><br>60 | 99 | 2.011 | -3.29 |
| *cys*G R | TTTGCCTTTTTGCGCTTC | Locus_Tag b3368 | | | |
| *M*1 F | GAAGCTGACCAGACAGTCGC | *M1* transcripts quantification (Ref decay assay (5))<br>377 | | | |
| *M*2 R | AGGTGAAACTGACCGATAAG | Locus_Tag b3123 | | | |

**References (1 to 5)**
